# A curvilinear coordinate flatmap for visualizing hippocampal structure and development

**DOI:** 10.64898/2026.01.29.702633

**Authors:** Ashwin A Bhandiwad, Fae Kronman, Josephine Liwang, Pan Gao, Cyrille Rossant, Dan Birman, Luis Puelles, Song-Lin Ding, Xiangmin Xu, Lydia Ng, Yongsoo Kim, Tyler Mollenkopf

## Abstract

The hippocampal formation is a highly curved and topographically complex forebrain structure. This complex geometry presents persistent challenges for analyzing subregional, laminar, and connectivity patterns. Here, we present a computational workflow that generates curvilinear-coordinate flatmaps from Common Coordinate Framework (CCF) registered hippocampal and retrohippocampal regions by solving the Laplacian equation to derive geodesic streamlines. This transformation unfolds the hippocampus into a planar slab, bounded by the meningeal and ventricular surfaces, with the depth defined along the radial axis. We apply this transform to image volumes, single neuron reconstructions, and point data, including spatial transcriptomic and rabies tracing datasets, revealing topographic variations in the dorsoventral and radial axes that are obscured in the CCF coordinate space. As proof of principle, we use flatmaps to show connectivity loss in a mouse model of Alzheimer’s disease and track postnatal development of microglial distribution in the hippocampus. This work provides an efficient and accessible resource for visualizing hippocampal organization across development and disease, offering new opportunities to interrogate the structure and function of this important brain region.

## Main

Topographic organization is a defining feature of neural circuits, linking spatial position to computation. In sensory and motor systems, topographic maps define how information is represented, integrated, and organized (Kaas, 1997). Understanding a region’s function therefore requires not only identifying cell types and their connections but also understanding their spatial and geometric context.

Advances in imaging, spatial transcriptomics, and single neuron morphological reconstruction have recently enabled the creation of large datasets that uncover cell type organization at a brain-wide scale (Kim et al., 2017; Liu et al., 2025; Wu et al., 2022; Yao et al., 2023). When integrated within a standardized atlas that share a common coordinate system, regional annotations, and consistent terminology, these datasets provide a powerful reference for understanding regional organization (Hawrylycz et al., 2023; Kleven, Gillespie, et al., 2023; Leergaard & Bjaalie, 2022).

Although whole-brain atlases exist for various species and developmental stages (Evans et al., 2012; Frey et al., 2011; Kleven, Bjerke, et al., 2023; Kronman et al., 2024; Liwang et al., 2025; Wang et al., 2020), the topographic organization of cell types and their connectivity remains difficult to assess. A key limitation to understanding topography is the use of a coordinate system with linear and orthogonal axes in the anterior-posterior, dorsal-ventral, and medial-lateral planes. While intuitive, this linear coordinate system fails to capture intrinsic curvature and nonlinear biological variation of brain structures, particularly across development (Bernhardt et al., 2022; Qiu et al., 2024; Thompson et al., 2014; Zhao et al., 2024). Therefore, resolving topographic organization requires a nonlinear coordinate system aligned with the primary axis of variation (Fig 1A).

**Figure 1:**
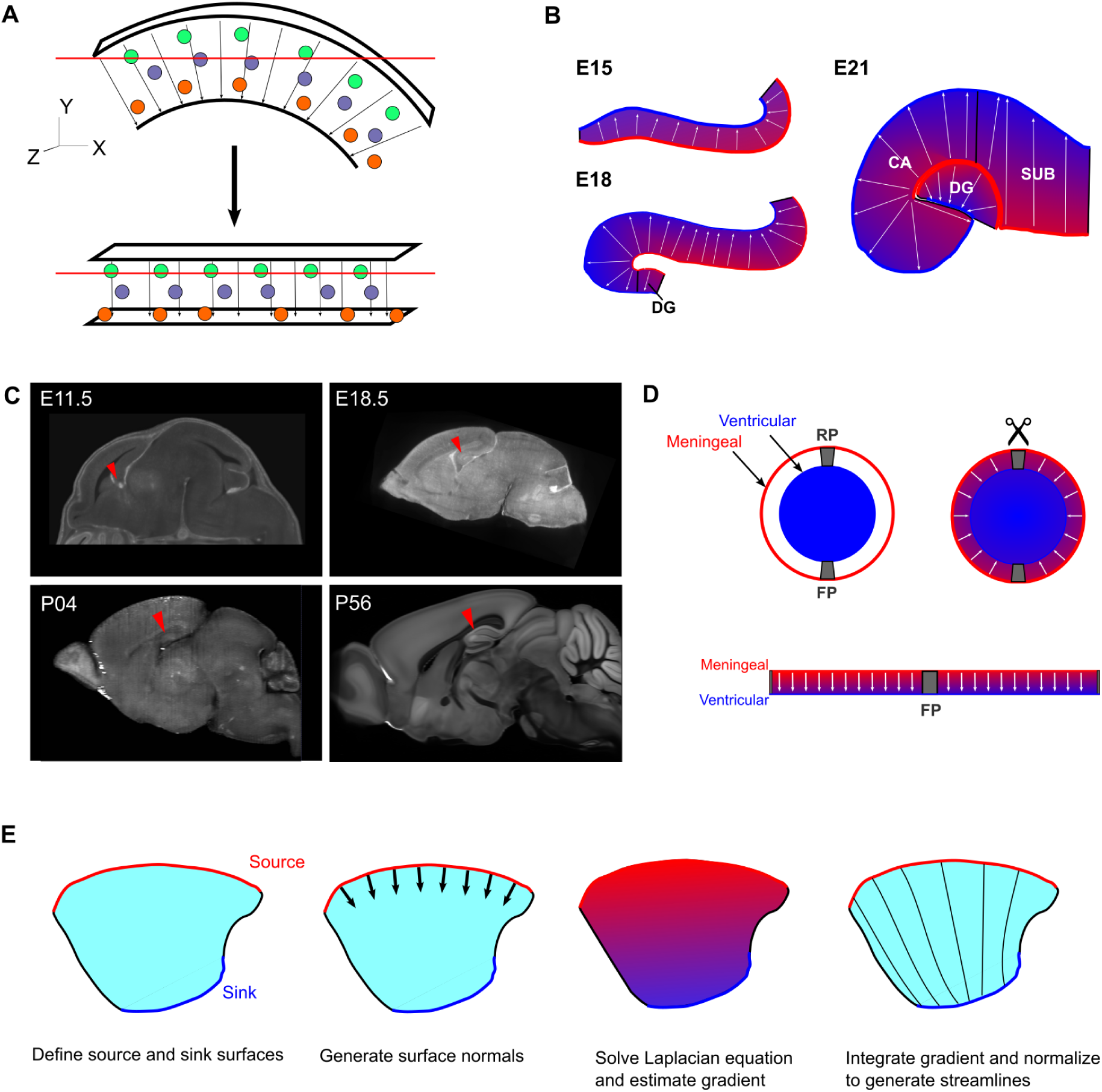
Conceptual framework for curvilinear-coordinate flatmapping of the hippocampus. **A,** Illustration of flatmapping to reveal topography of cell types in a volume. Traditional sectioning along CCF (x,y, z) axes (top) captures cells at multiple depths. Transformation to curvilinear coordinates (bottom) parameterizes the volume so that an equivalent section captures cell topography along the radial axis. **B,** Illustration of developmental progression of hippocampus at E15, E18, and E21 with meningeal (red) and ventricular (blue) surfaces annotated in sagittal views. Arrows indicate streamline orientation and shading shows the Laplacian gradient. **C,** Sagittal views of serial two-photon templates of E11.5, E18.5, P04, and P56 showing hippocampal development. Arrowheads show hippocampal formation at each developmental stage. **D,** Definition of outer (meningeal) and inner (ventricular) surfaces based on the neural tube model, conceptualized as a slab with the radial dimension representing cortical depth. **E,** Overview of streamline generation. Surface normals define boundary conditions for Laplace’s equation, and integration of the resulting gradient field produces smooth trajectories between the two surfaces.

Geodesic flatmaps address this limitation by unfolding structures onto simpler manifolds while preserving intrinsic geometry (Fischl et al., 1999; Hahn et al., 2021; Hahn & Duckworth, 2023). Early implementations were qualitative and communicated three-dimensional data on paper, a two-dimensional medium (Sherk, 1992; Van Essen & Maunsell, 1980). More recently, quantitative flatmaps, created using streamlines, meshes, and physical models (Bolaños-Puchet et al., 2024; DeKraker et al., 2018; Diers et al., 2023; Jones et al., 2000; Ng et al., 2010; Swanson & Hahn, 2020; Wu et al., 2022), provide a data-driven approach to understanding a structure’s nonlinear geometry. Streamline-based flatmaps have been applied to visualize connectivity, density, and vascular organization in mouse cortex (Bennett et al., 2024; Harris et al., 2019; Peng et al., 2021; Wu et al., 2022).

The hippocampal formation (HPF), a key structure for learning, memory, and emotional control (Knierim, 2015), poses a unique challenge for flatmapping because of its high curvature. Existing hippocampal flatmaps remain largely qualitative without computational means to apply them to data assets (Swanson & Hahn, 2020). Here, we introduce a resource for transforming hippocampal volumes into a curvilinear coordinate system that remains topologically consistent across development. Using image volumes from the Allen CCFv3 mouse atlas and early postnatal development atlases (Liwang et al., 2025; Wang et al., 2020), we define geodesic hippocampal coordinate spaces for adult (P56) HPF and two developmental stages, postnatal day (P)4 and P14. This framework preserves laminar and longitudinal organization while providing a common space for integrating multimodal data from mesoscale connectivity, single-neuron reconstructions, and spatial transcriptomic datasets. The resulting flatmaps reveal hippocampal organization in an intuitive coordinate system, offering an accessible tool for exploring hippocampal topography and its developmental origins.

## Results

To generate developmentally consistent flatmaps of the hippocampal formation (HPF), we began with its embryonic origin. The HPF begins as a sheet derived from the cortical hem and curves into its characteristic crescent shape by embryonic day 18 (E18) (Angevine, 1975; Bayer, 1980; Swanson & Hahn, 2020). By E21, the dentate gyrus emerges as a distinct cap at the end of the cornu ammonis (CA)(Fig 1B). During this process, the original sheet differentiates into outer (meningeal) and inner (ventricular) surfaces that remain topologically consistent across development (Fig 1B-C). Based on the prosomeric model(Puelles et al., 2013), we aligned slab depth along the radial dimension with meningeal and ventricular surfaces as boundaries (Fig 1D-E). This framework provides a common geodesic space for comparing hippocampal organization across development.

Based on this scaffold, we implemented a curvilinear coordinate framework based on the streamline-based coordinate framework for the isocortex (Wang et al., 2020). We defined the meningeal and ventricular boundaries of the adult mouse hippocampal formation (HPF) using anatomical annotations from the Allen Mouse Brain CCFv3 (Wang et al., 2020). HPF subregions (CA fields, subiculum, presubiculum, postsubiculum, parasubiculum, and entorhinal cortex) and the dentate gyrus (DG) were isolated, with the DG processed separately because of its developmental discontinuity. Surface voxels were manually classified as meningeal, ventricular, or unassigned (Fig 1E, Fig 2A-B) to serve as boundary conditions for constructing a curvilinear coordinate system.

**Figure 2:**
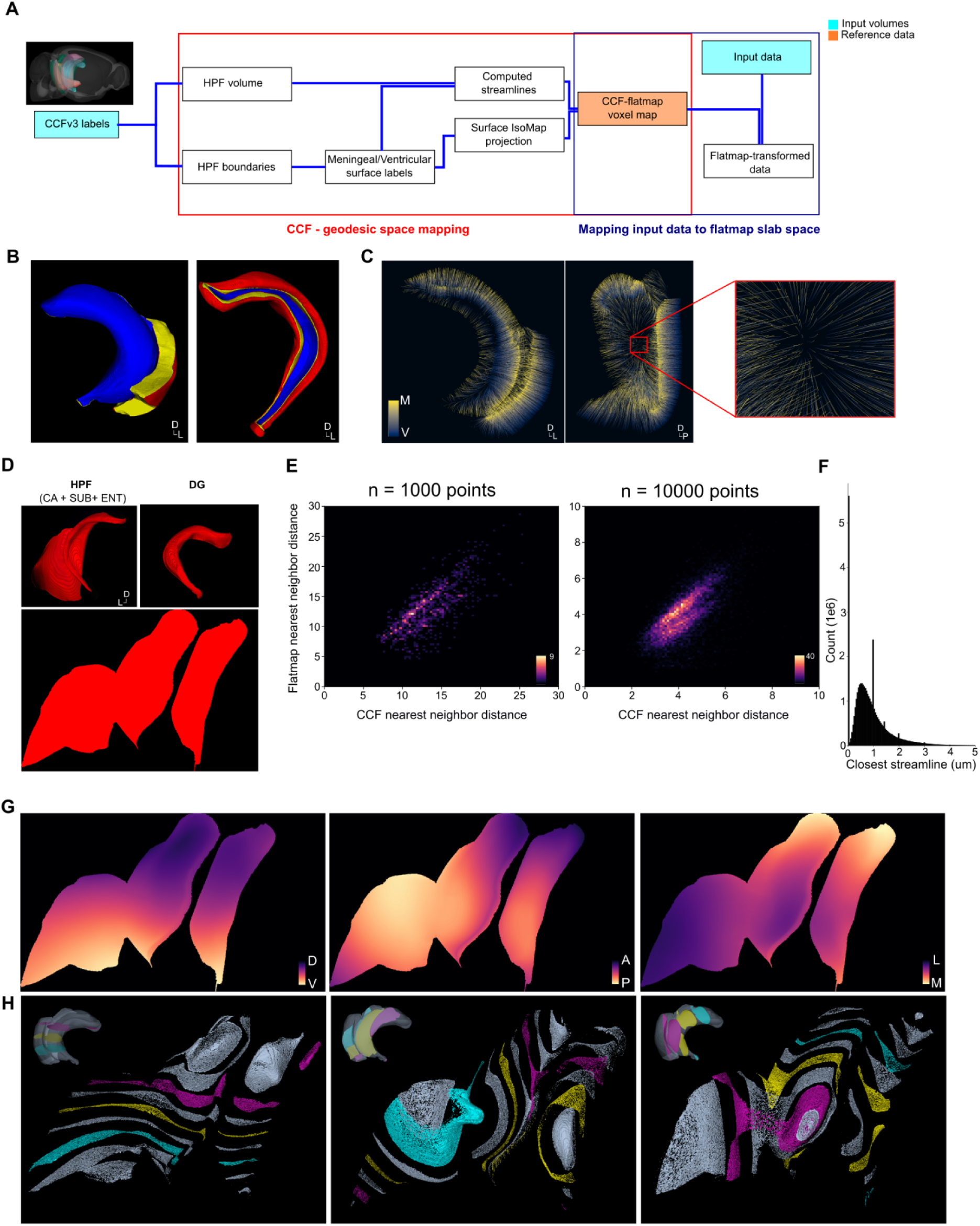
Workflow and flatmap creation for young adult (P56) HPF and DG. **A,** Overview of flatmapping workflow for CCF-registered volumes. Annotations and input volumes are processed to generate reference data assets which can be applied to subsequent image volumes. **B,** 3D reconstructions of coronal view of P56 HPF (left) and DG (right) volumes labeled as either meningeal-derived (red), ventricular-derived (blue) or edge (yellow) surfaces. **C,** Example streamlines through HPF volume colored by potential gradient in coronal (left) and sagittal (right) views. Inset shows zoomed view showing local streamline curvature. **D,** Isomap projection of HPF and DG meningeal surfaces. Top shows surfaces in CCF space. Bottom shows 2D flattened surfaces with manually aligned HPF and DG used for visualization. **E,** Histogram showing relationship between subsampled points in HPF and their nearest neighbor at n = 1000 points (left) and n = 10000 points (right), colored by voxel count. **F,** Histogram of closest streamline distance in CCF space for each voxel. **G,** Gradients of dorsal–ventral, anterior–posterior, and medial–lateral axes in flatmap space. Maps show midthickness section and colormaps show direction of gradient. **H,** 3D reconstructions of 200-µm sections along each axis in flatmap space, illustrating nonlinear warping of anterior–posterior and medial–lateral axes versus preserved dorsal–ventral alignment.

We solved Laplace’s equation between the meningeal and ventricular surfaces to obtain a smooth harmonic potential and integrated streamlines through this field to generate unique, non-intersecting trajectories linking the two surfaces (Fig 2C). Each voxel was assigned to a single streamline, defining a continuous mapping between CCF and curvilinear coordinates.

To obtain a two-dimensional embedding, we projected the meningeal surface into a geodesic slab using Isomap, which preserves local structure by computing shortest paths along a neighborhood graph(Tenenbaum et al., 2000) (Fig 2D). Nearest-neighbor correlations between CCF and streamline coordinates confirmed preservation of local spatial relationships (HPF: r = 0.70; DG: r = 0.17; Fig 2E). Although the embedding introduced some compression due to streamline downsampling, local spatial relationships remained well preserved (Fig. 2F).

Flattening a convex structure to a planar slab introduces either distance or volume distortion (Milnor, 1969). We quantified local volumetric distortion by calculating the local Jacobian determinant of the transformation. Compression was most pronounced in ventral CA, particularly near the ventricular surface, and more evenly distributed near the meningeal surface (Fig S1), indicating that volumetric inferences across subregions should be interpreted with caution.

To assess biological interpretability, we examined how CCF (Cartesian x,y,z) axes were transformed into flatmap space. Gradients along the dorsal–ventral, anterior–posterior, and medial–lateral axes were mapped through the combined HPF and DG mask. The dorsal–ventral axis was largely preserved, whereas the other two axes underwent nonlinear transformations across the slab (Fig 2G-H). Given the well-established functional and molecular gradients along the dorsal-ventral axis (also known as the septal-temporal or longitudinal axis)(Fanselow & Dong, 2010), we oriented flatmaps with the vertical axis aligned approximately to the CCF dorsal-ventral axis. This orientation helps highlight features along a well-understood and biologically relevant dimension.

### Application to mesoscale anterograde and single neuron reconstruction data

We next tested whether the curvilinear coordinate space captures known spatial connectivity patterns. Average template and CCFv3 structure labels (Wang et al., 2020) were first transformed into flatmap space (Fig 3A).The hippocampal formation unfolded into a geodesic planar slab with well-preserved laminar organization.

**Figure 3:**
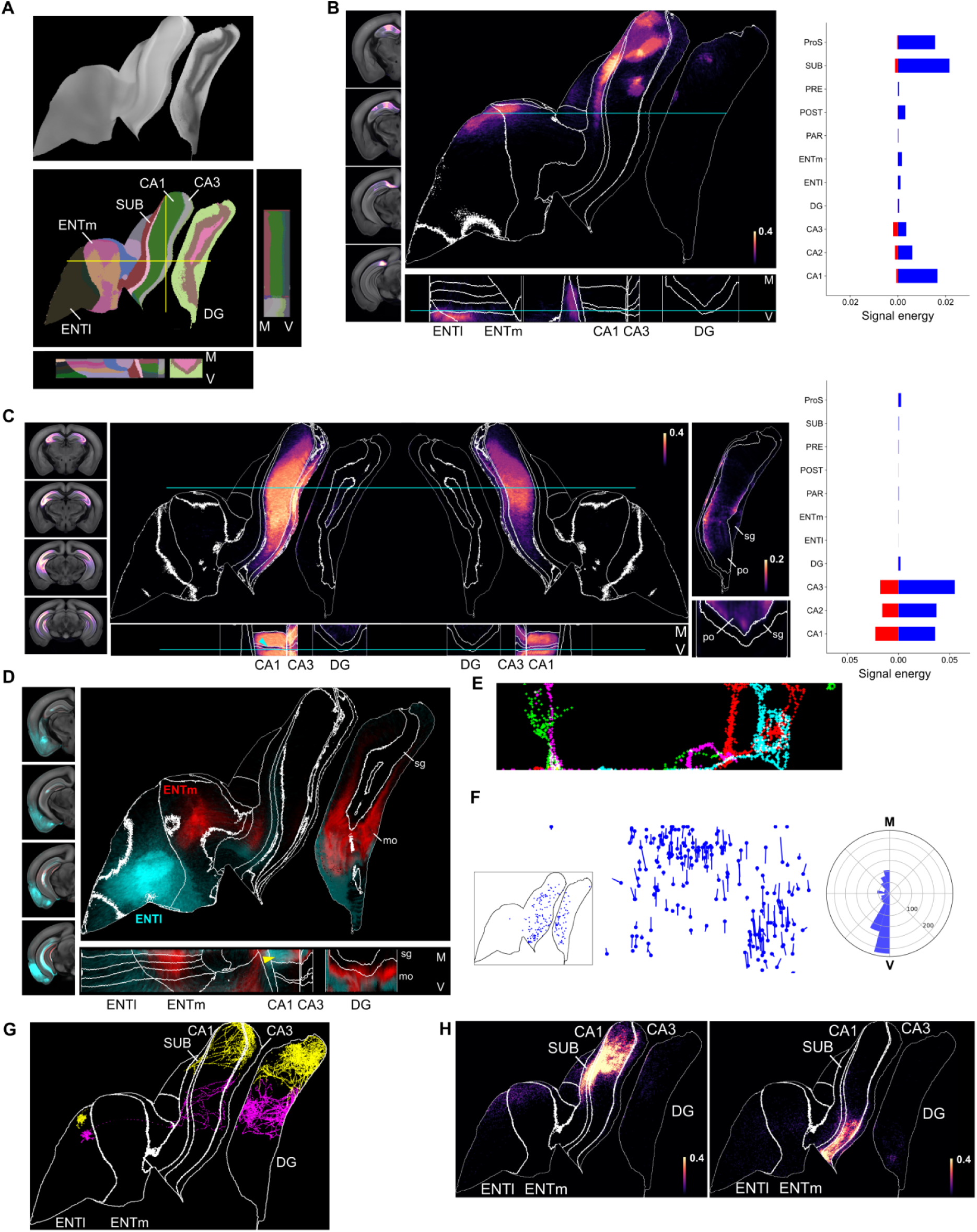
Flatmaps of mesoscale and single neuron connectivity in the hippocampal formation. **A,** Midthickness flatmap of the STPT average template with matching CCFv3 annotation, showing continuous laminar organization. Lines indicate sections for orthogonal view on bottom and right. CA1: Cornu Ammonis 1, CA3: Cornu Ammonis 3, DG: Dentate gyrus, ENTm: Medial entorhinal cortex, ENTl: Lateral entorhinal cortex, SUB: Subiculum. **B,** Averaged signal intensity for injections into CA1 (n = 8 experiments). Right, signal energy (summed voxel intensity normalized by number of voxels) by region. Blue shows ipsilateral signal, red shows contralateral signal. **C,** Left: Averaged signal energy for injections into CA3 (n = 7 experiments) and their topography in a bilateral flatmap. Blue arrowhead shows lack of signal in the CA1 pyramidal layer. DG cutout on the right shows the CA3 signal with polymorph (po) and granule cell (sg) layers annotated. Right: signal energy in each subregion within HPF and DG showing high signal in ipsilateral (blue) and contralateral (red) CA1 and CA3. **D,** Midthickness flatmap section showing averaged mesoscale injections into lateral (cyan) and medial (red) entorhinal cortices and their topographic separation in DG. Bottom shows laminar distribution of ENTl and ENTm projections within DG. Arrowhead shows the CA1 stratum lacunosum moleculare (slm). **E,** Example single neuron reconstructions in flatmap space. **F,** Soma positions and initial axon orientation vectors from sampled HPF neurons (n = 1006). Right: Circular histogram showing heading direction along meningeal-ventricular axis. **G,** Example L1 ENTm (yellow) and L2/3 ENTm (magenta) neurons in HPF with distinct projection zones in CA1 and DG. **H,** Putative axon terminal distribution for single neurons sampled from dorsal (left) and ventral (right) CA1. White dots show soma locations.

We then applied the transformation to mesoscale anterograde connectivity data from the Allen Mouse Connectivity Atlas (Oh et al., 2014). Hippocampal connectivity exhibits characteristic topographic gradients along the dorsoventral and transverse axes (Fanselow & Dong, 2010; Strange et al., 2014; Witter et al., 1989), providing a benchmark for validating the flatmap coordinate space. Aggregated CA1 injection experiments (n = 8) revealed dense projections to the subiculum, prosubiculum, and in layers 5-6 of the entorhinal cortex, consistent with previous work (Amaral & Witter, 1989; Cenquizca & Swanson, 2007)(Fig 3B). CA3 injections (n = 7 experiments) showed ipsilateral and contralateral projections to CA1, contralateral CA3, and DG polymorph layer, corroborating previous work (Ishizuka et al., 1990; Li et al., 1994; Swanson et al., 1978)(Fig 3C). Notably, coordinate transformation preserved the relative absence of signal in the CA1 pyramidal layer, underscoring the accuracy of the transformation in maintaining anatomical detail.

We then evaluated whether flatmaps captured laminar specificity of entorhinal inputs to DG. Projections from the lateral entorhinal cortex (ENTl) terminate in the outer DG stratum moleculare, while medial entorhinal cortex (ENTm) projections terminate in the inner DG(van Groen et al., 2003).Averaged ENTl and ENTm projections localized to the expected layers after transformation (Fig 3D). Laminar segregation was also present in the CA1 stratum lacunosum moleculare (slm), demonstrating that the transformation preserves laminar specificity alongside broader topographic structure.

Because mesoscale data aggregates fibers from multiple layers, cell types, and includes fibers of passage, we extended the curvilinear transformation to single neuron reconstructions. We transformed reconstructions from the Janelia MouseLight browser (Chandrashekar, 2017) and the mouse projectome atlas (Qiu et al., 2024), and selected 10,158 neurons with soma within the HPF. In flatmap coordinates, axons were aligned vertically along the meningeal-ventricular axis, reflecting radial geometric organization (Fig 3E-F). Two example neurons from the lateral entorhinal cortex - one in layer 1 and one in layers 2/3 - showed distinct projection zones in dorsal DG (Fig 3G). At the population level, CA1 outputs also showed dorsoventral gradients. Dorsal CA1 neurons targeted subiculum and dorsal entorhinal cortex, whereas ventral CA1 neurons projected to ventral targets (Fig 3H).

Together, these analyses show that curvilinear coordinates preserve laminar fidelity while revealing large-scale topographies in both mesoscale and single-cell datasets. Flatmaps therefore provide a unified framework for visualizing hippocampal connectivity across scales.

### Topographic organization of cell types in flatmap coordinate space

We next tested whether flatmaps could resolve the spatial organization of molecularly defined cell types. We transformed a registered spatial transcriptomic dataset comprising 812,084 HPF cells from multiple sources (Yao et al., 2023; Zhang et al., 2023), each mapped to a cell type ontology. As a first step, we transformed cells labeled by putative neurotransmitter identity, which showed uniform distribution across the HPF (Fig 4A).

**Figure 4:**
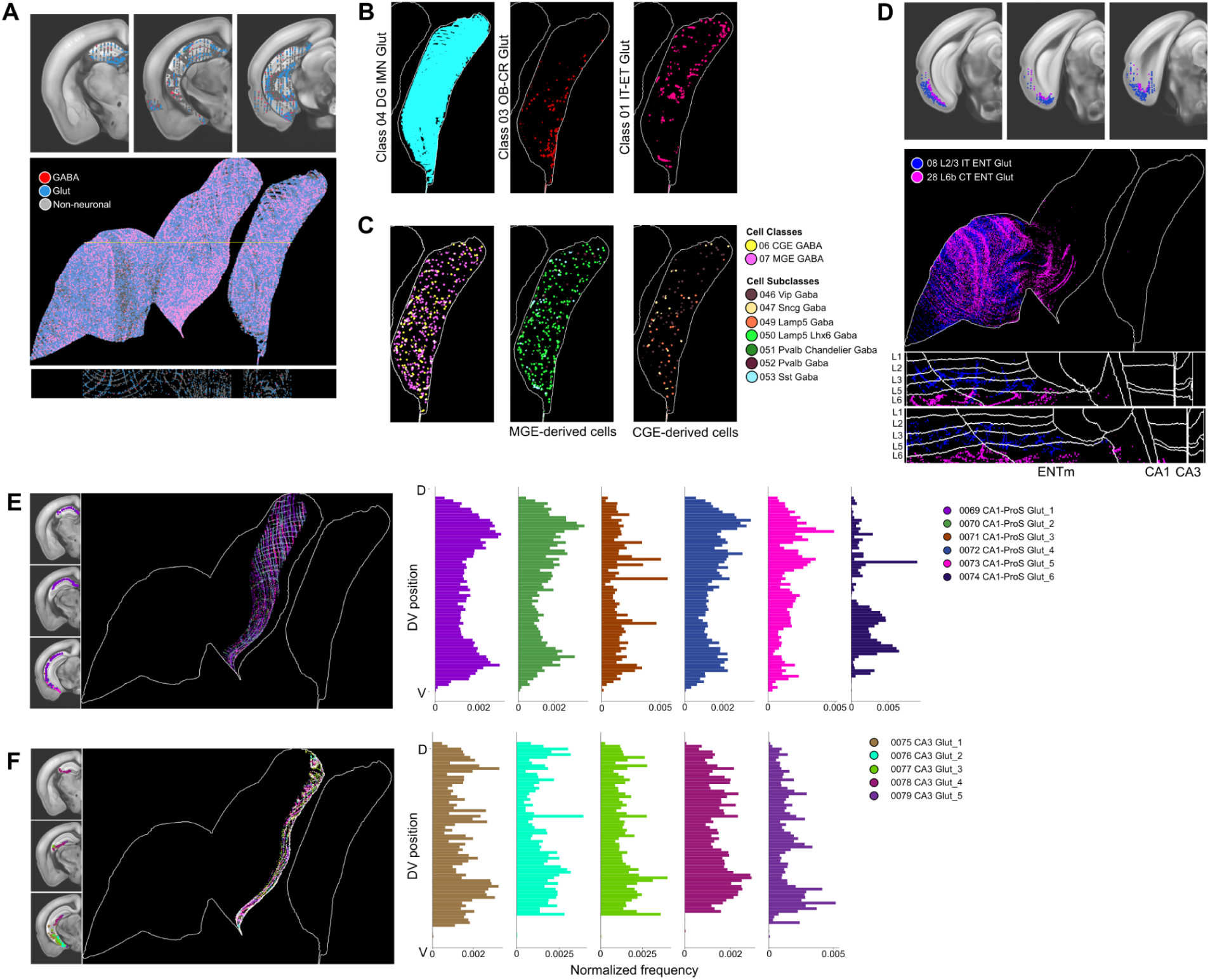
Spatial transcriptomic topography visualized in curvilinear flatmap space. **A,** Cells from a registered spatial transcriptomic dataset (n=812,084) colored by neurotransmitter identity showing consistent laminar organization across HPF. Top shows cells in CCF space, bottom shows maximum projection of cells in flatmap space. **B,** Maximum projection of glutamatergic cells in the dentate gyrus granule layer labeled by cell class. **C,** Maximum projection flatmap of GABAergic cells. Left: GABAergic cells labeled by cell class. Maximum projection of MGE GABA cells (Middle) and CGE GABA cells (Right) in the dentate gyrus granule cell layer labeled by subclass. **D,** Maximum projection flatmap of L2/3 IT ENT Glut (magenta) and L6b CT ENT Glut (blue) subtypes. Bottom shows orthogonal views with regional and laminar boundaries overlaid. **E,** Maximum projection of topography of CA1-ProS Glut supertypes in CA1 pyramidal layer. Cells are colored by supertype identity. Left: Coronal series showing supertype distribution in CCF space. Right: Histograms of cell distribution along dorsal-ventral axis split and labeled by CA1-ProsS Glut supertypes. **E,** Maximum projection of topography of CA3 Glut supertypes in CA3 pyramidal layer. Cells are colored by supertype identity. Right: Histograms of cell distribution along dorsal-ventral axis split and labeled by CA3 Glut supertypes

To examine topographic organization of glutamatergic and GABAergic cell subtypes, we focused on the dentate gyrus granule cell layer (DGsg), which contains the primary excitatory cell type within the dentate gyrus. Most glutamatergic cells were immature DG (DG-IMN Glut) granule cells, but two additional classes showed striking positional biases: Olfactory bulb-Cajal Retzius (OB-CR Glut) cells were enriched ventrally, while long-range projecting glutamatergic cells localized dorsally (Fig 4B). Among GABAergic cells, medial ganglionic eminence-derived classes dominated overall, with Vip GABA cells enriched dorsally and caudal ganglionic eminence derived Lamp5 GABA cells confined ventrally (Fig 4C). This pattern suggests that inhibitory interneuron distribution reflects developmental lineage along the dorsoventral axis.

Next, we focused on cell type laminar distribution in the entorhinal cortex along the meningeal-ventricular axis. Corticothalamic layer 6b (L6b CT ENT Glut) cells mapped as expected to deep layers 5–6, whereas intratelencephalic layer 2/3 (L2/3 IT ENT Glut) cells were present in superficial layers but unexpectedly were also present in deeper ventral ENTm (Fig 4D). Flatmaps thus enable visualization of subtle laminar deviations that are difficult to discern in CCF coordinate space.

Finally, we assessed supertype-level organization of CA1 and CA3 pyramidal neurons. CA1-ProS Glut cells (six supertypes) showed a bimodal distribution along the dorsoventral axis, with the rare CA1-ProS Glut 6 subtype enriched ventrally (Fig. 4E). In contrast, CA3 Glut neurons were more evenly distributed, with a modest bias toward ventral CA3 (Fig. 4F). Together, these analyses demonstrate that curvilinear transformation captures both laminar and topographic organization of cell types across multiple levels of the ontology, linking transcriptomic identity to spatial patterning within the HPF.

### Application of flatmaps to retrograde labeling data in a model of Alzheimer’s disease

To test whether flatmaps can reveal spatial features of pathology, we applied the coordinate transformation to retrograde tracing data from 5xFAD mice, a transgenic model of Alzheimer’s disease. 5xFAD and wild-type (WT; C57BL/6J) littermates received injects in the dorsal CA1 with AAV helper viruses (AAV1-CamKII0.4.Cre.SV40 and AAV8-DIO-TC66T-2A-GFP). Seventeen days later, EnvA-SADΔG-tdTomato was injected in the same coordinates, and mice were perfused 9 days post-rabies injection, at approximately 4.3 months of age.

In WT animals, monosynaptic rabies virus mapped input neurons were detected in CA1, CA3, subicular complex, and entorhinal cortex (Fig. 5A). Flatmap transformation revealed a laminar arrangement of these inputs along the meningeal–ventricular axis, aligned with starter cell distribution. This organization was not apparent in CCF space.

**Figure 5:**
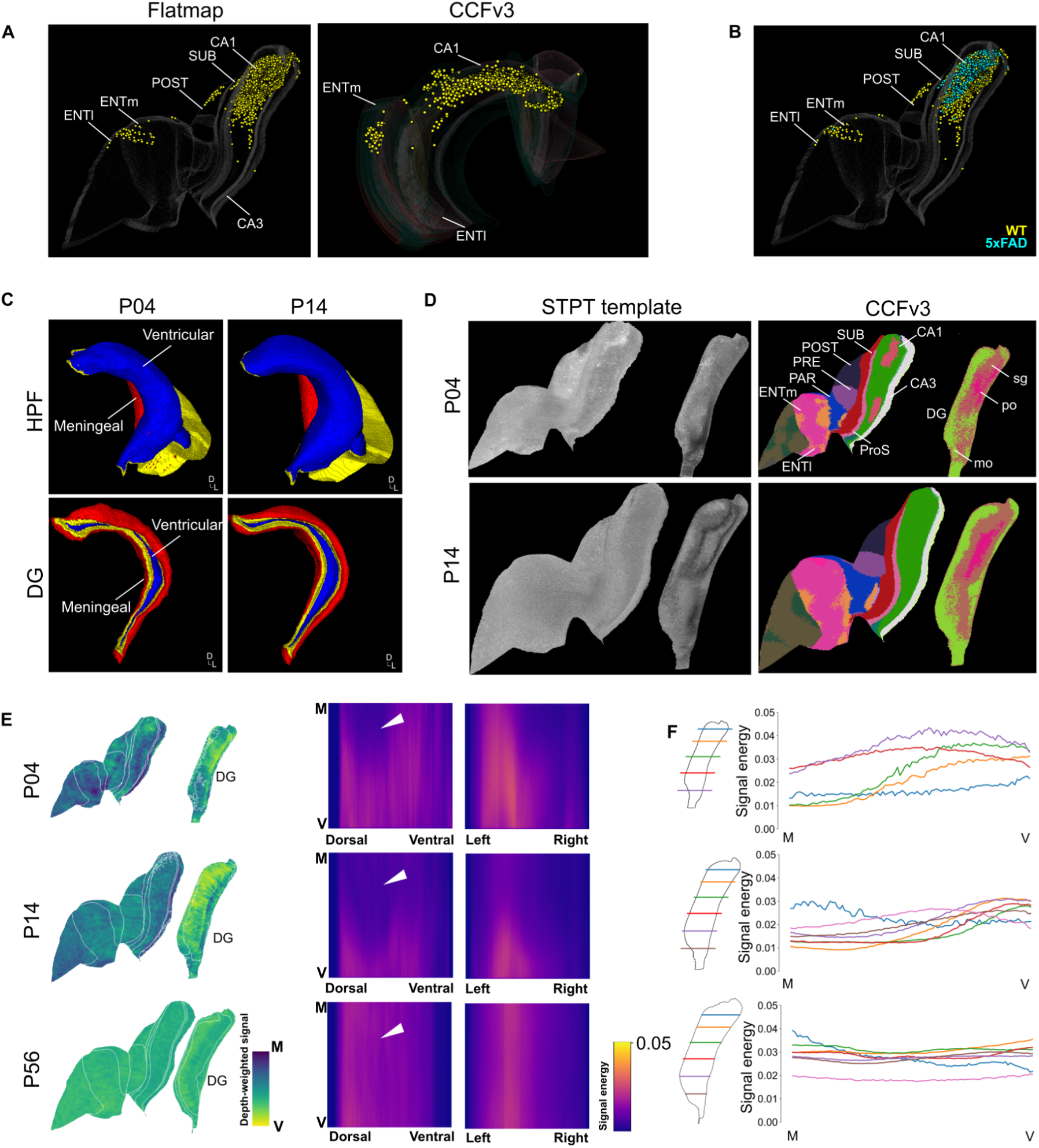
Applying curvilinear-coordinate flatmaps to disease and developmental analysis. **A,** Retrograde labeling in WT and 5xFAD mice shown in flatmap (left) and CCF space (right). Presynaptic cells are labeled in yellow, starter cells are labeled in magenta with HPF outline (white) as a reference. **B,** 3D view of presynaptic cell distribution in HPF flatmap coordinate space between WT (yellow) and 5xFAD (cyan) animals. Starter cells for WT (n=17) and 5xFAD (n=14) are not shown. **C,** 3D reconstructions of coronal view of P04 (left) and P14 (right) surfaces labeled as either meningeal (red), ventricular (blue) or edge (yellow). Top shows HPF and bottom shows DG. **D,** Midthickness sections for STPT template flatmaps in P04 (top) and P14 (bottom) with associated CCFv3 flatmapped labels. **E,** Depth-weighted flatmaps of P04 (top), P14 (middle), and P56 (bottom) showing microglia topography labeled by Cx3cr1-GFP. Colormap shows signal density along the meningeal-ventricular axis. Right, cross-sectional views of Cx3cr1-GFP signal at sections defined by horizontal black lines. Colormap shows signal intensity values. **F,** Quantification of average Cx3cr1-GFP signal intensity along the DG meningeal-ventricular axis quantifies axis-specific changes in microglial distribution over development.

Comparing WT and 5xFAD mice with matched starter cell numbers (n=17 WT, n=14 5xFAD)(Fig 5B, S3) showed that both groups preserved laminar organization, but 5xFAD mice displayed fewer presynaptic neurons overall, especially in the entorhinal cortex and subiculum. Presynaptic CA1 neurons were also reduced and occupied a shorter span along the hippocampal longitudinal axis.

These findings demonstrate that coordinate transformation can capture subtle spatial changes in connectivity and propose flatmaps as a practical tool for visualizing differences in cell distribution and connectivity in models of neurodegeneration.

### Curvilinear coordinate space maps for spatiotemporal analyses across development

Finally, we asked whether flatmaps could track hippocampal topography across development. Early postnatal development involves inhibitory neuron migration, synaptogenesis, and changes in neuronal activity that shape hippocampal function (Cossart & Khazipov, 2022; Garaschuk et al., 1998; Olsen et al., 2023; Wong et al., 2005). Because our model is defined based on meningeal and ventricular boundaries, we expected curvilinear coordinates to yield a developmentally invariant latent space with preserved topology.

Annotated surfaces were registered sequentially from P56 to P14 and P4 and manually refined to correct minor registration errors. Curvilinear coordinates were computed independently for HPF and DG at each stage, and corresponding CCFv3 annotations were transformed for comparison and labeling (Fig 5C). Despite differences in the shape and size of the HPF between P4 and P14, CCFv3 labels maintained a consistent structure, suggesting feasibility for cross-development analyses (Fig 5D). However, subregion volumes showed nonlinear distortions between CCF and flatmap space. CA pyramidal layers preserved relative proportions, but regions with high curvature (eg. ENTm, DG, prosubiculum) diverged across development (Fig S2). These distortions emphasize that distance-based rather than volumetric measures provide more reliable developmental comparisons.

To test whether flatmaps capture developmental dynamics, we transformed published Cx3cr1-eGFP volumes, which label microglia, at P4, P14, and P56(Liwang et al., 2025). Microglia, which regulate synapse formation and cell death, proliferate and colonize brain regions with axis-specific density differences during early postnatal development (Paolicelli et al., 2011; Barry-Carroll & Gomez-Nicola, 2024). At P4, expression was enriched in ventral DG (ventricular side) and dorsal DG (meningeal side) (Fig 5E-F). By P14, Cx3cr1 expression was more uniform dorsoventrally but shifted towards the meningeal surface in the ventral DG, consistent with radial migration. In the entorhinal cortex, Cx3cr1 expression increased from the ventricular to meningeal surfaces between P4 and P14. By P56, expression was uniformly distributed across the HPF, consistent with prior observations (Barry-Carroll et al., 2023; Jinno et al., 2007). These axis-biased shifts recapitulate known microglial migration and colonization patterns, demonstrating the value of flatmaps for developmental analysis.

## Discussion

In this work, we present a curvilinear coordinate system that “unfolds” the hippocampal and retrohippocampal formation into a planar slab with preserved nearest-neighbor distances. Using multiple data modalities, we show that this approach captures topographic variation that is obscured in CCF coordinates. While hippocampal organization across the longitudinal axis is well defined, this coordinate space extends analysis to the radial axis in a geodesic space. The resulting “2.5D” representation captures both surface and depth-related variation.

The data assets and transformation scripts are available at https://github.com/AllenInstitute/flatmap. Using this resource, a 10 µm resolution volume can be transformed on a local workstation in minutes, making it accessible for a wide range of research applications.

Our model is based on a radial organization inspired by the concentric ring model of development (Puelles et al., 2019). Anchoring to meningeal and ventricular surfaces yields a developmentally conserved axis that supports comparison across postnatal development. This approach also makes the workflow extensible. Flatmaps could be used to align hippocampal structures across species using streamline-derived coordinates (DeKraker et al., 2018) or compare within-species comparisons, eg. between HPF and isocortex.

A key advantage of transforming data into a geodesic space is its ability to reveal spatial details that are not apparent in CCF space. For example, mesoscale connectivity revealed laminar segregation of DG inputs at the ENTm-ENTl boundary, a pattern consistent with previous observations but now quantifiable from volumetric data (Fig 3D). Likewise, developmental flatmaps captured microglial migration along the meningeal–ventricular axis, consistent with histological reports of colonization gradients. These correspondences indicate that the transformation preserves biological signals and highlight its utility for hypothesis generation, eg. testing whether dendritic targeting by ENT inputs is systematically coupled to dorsoventral cell type variation.

One limitation of this approach is area and volume distortion, particularly in developmental flatmaps (Fig S1, 5D). Distortions arise from redundancy and downsampling when mapping to streamlines, particularly in smaller structures or those with high curvature, such as the ventral CA and meningeal DG. While distance-based analyses are robust, volumetric inferences require caution. Because distortions accumulate at larger volumes, applications where volumetric precision is required may require alternative approaches, such as mesh-based transforms.(Bolaños-Puchet et al., 2023)

Boundary artifacts are another challenge, particularly at meningeal and ventricular edges and the CA3–DG interface (Fig 4G). Mis-mapping can obscure critical pathways such as the perforant path, an important connectivity pathway between the entorhinal cortex and the DG,(Amaral & Witter, 1989) and fragment single neuron morphologies when axons exit the HPF boundary. These artifacts highlight the need for refinement of surface definitions and expansion of this approach to more brain regions to preserve continuity. Addressing these boundary-related artifacts will be crucial for improving the accuracy of structural mapping, particularly in developmental and connectivity studies.

An important question moving forward is how flatmaps generalize across data modalities. This study highlights flatmap transformation for point data (e.g., MERFISH soma) and continuous fields (e.g., mesoscale connectivity). These data are differentially affected by distortions. Systematic evaluation of these differences, and integration with ground-truth histology, will be essential for defining reliable use cases of the transform.

Despite these caveats, the approach is highly generalizable. Because it requires only two boundary surfaces, it can be applied across species and to other folded structures in the brain and body. The method also integrates with emerging spatial omics approaches such as MERFISH (Hu et al., 2021; Townes & Engelhardt, 2023). Application of curvilinear coordinate transformation to spatial single cell data also offers the opportunity to link spatial context to cell identity, connectivity, and dynamic processes like development and disease. Flatmaps thus provide a computational framework for analyzing topologically complex structures, enabling comparative and developmental analyses that are otherwise obscured in CCF coordinate space.

## Methods

### Data sources

The average template and annotations for the adult P56 brain were published previously (Wang et al., 2020) and accessed through the Allen software development kit (Allen SDK). Mesoscale anterograde tracing data were accessed through the Allen Mouse Brain Connectivity portal (https://connectivity.brain-map.org) and the MERFISH spatial transcriptomic data was accessed through the Allen Brain Cell (ABC) atlas webpage (https://portal.brain-map.org/atlases-and-data/bkp/abc-atlas). Single neuron reconstructions were accessed from the Janelia MouseLight browser (https://ml-neuronbrowser.janelia.org) and the Mouse Projectome Atlas (https://mouse.digital-brain.cn/projectome/hipp). All neurons with soma in the HPF were selected for transformation.

Developmental templates and annotations for postnatal stages P4-P14 were published previously (Liwang et al., 2025). CCFv3 annotations were mapped onto the P4 template by nonlinear registration using Advanced Normalization Tools (ANTs)(Avants et al., 2011) using the AntsPy python library (https://github.com/ANTsX/ANTsPy/).

### Meningeal and ventricular surface annotation

Meningeal and ventricular surfaces were annotated manually using ImageJ. Each annotation volume was masked to generate a volume for the hippocampal formation (HPF). The hippocampal formation was split into the dentate gyrus and a combined structure of the Ammon’s horn, subiculum, presubiculum, postsubiculum, parasubiculum, and entorhinal cortex. The dentate gyrus is discontinuous at these developmental stages and was processed as a separate volume.

The annotation volume was filtered to include all descendants of the HPF (CA, PRE, SUB, ProS, PARA) and DG in the CCFv3 ontology (Wang et al., 2020). The filtered volume was binarized to generate a region mask and the edges of the region mask were used as the bounding surface of the region of interest. Because the CCFv3 volume is symmetric, the following steps were performed on a single hemisphere to minimize computational costs.

Meningeal and ventricular surfaces were manually defined using pixel painting. For each section, edges were compared to the surfaces in the filtered annotation and annotated as either meningeal, ventricular, or unassigned. The meningeal surface was defined as the surface derived from the meningeal layer (Fig 2B). Where possible, the ventricular layer (eg. L6 of entorhinal cortices, CA stratum oriens) was used as the ventricular surface. For the dentate gyrus, meningeal and ventricular boundaries were extended from the contiguous regions in CA3. Unassigned (“edge”) surfaces were defined as any surface that was neither meningeal nor ventricular. In hippocampal regions with no laminar subdivisions, the known laminar meningeal/ventricular boundaries were extended from layered structures by visual inspection to fill the continuous plane.

### Streamline computation

To find the shortest-distance path through the curved volume, the protocol used streamlines derived from a Laplacian gradient (Fig 2A). Theory and details of streamline generation have been published previously (Jones et al., 2000; Ng et al., 2010; Wang et al., 2020). Streamline computation was performed using a forked implementation of the IBL atlas project (https://github.com/int-brain-lab/atlas). Streamlines were generated from the solution to Laplace’s equation. Briefly, the annotated edges and region volume combined and length-normalized orthogonal vectors to the meningeal surface were computed (Fig 1E). Laplace’s equation was defined with Dirichlet boundary conditions at the meningeal surface and Neumann boundary conditions at the ventricular and edge surfaces. The partial differential equation was solved using numerical approximation using an iterative procedure in a GPU implementation. Following empirical data from cortical streamline generation, n=10000 iterations were used to arrive at convergence. The solution to Laplace’s equation was used to estimate a gradient from the meningeal to ventricular surfaces in a stepwise process with forward and backward differences depending on whether the voxel was within the volume or on the bounding surfaces.

Curvilinear streamlines were computed by integrating the Laplacian gradient using an ordinary differential equation. Gradients were integrated from meningeal to ventricular surfaces using the Euler method. Finally, streamlines were rescaled so that each streamline consisted of n=100 points sampled evenly along the meningeal-ventricular axis.

### Mapping CCF to curvilinear (flatmap) slab

To generate the flattened slab space from streamlines, a 2D manifold of the 3D meningeal surface was generated using Isomap (Tenenbaum et al., 2000). Isomap is a dimension reduction technique that preserves local nearest-neighbor geodesic (ie. shortest path) distances along a surface. Isomap was performed using python’s SciPy library. A local neighborhood of n=25 nearest neighbors was empirically determined as the neighborhood that preserved distances and balanced against high spatial clustering.

To reconstruct streamlines, the meningeal surface was subsampled to 120,000 voxels, preserving its shape while reducing memory load and computational costs. In cases where the meningeal surface was significantly smaller than the ventricular surface, such as in the dentate gyrus, the origin of the streamlines was inverted with the ventricular surface assigned as the streamline origin. This adjustment improved the representation of the volume because the source surface had a larger surface area than the sink - this reduced compression artifacts, particularly near the edges. After integrating these adjustments, the flatmap space was represented as the Isomap flattened meningeal surface as the (x,y) coordinate and streamline depth (n=100 steps) as the depth coordinate (Fig 2D).

Bilateral planar slabs were generated by orienting the HPF and DG surfaces along primarily the dorsoventral axis (Fig 2G). Because HPF and DG were independently processed, surfaces were manually aligned so that the distal CA3 boundary and proximal DG boundary were roughly aligned. Moreover, independent Isomap transformation and streamline generation led to an overestimation of the DG surface, which was corrected by scaling the DG surface by the ratio of HPF to DG meningeal surface areas in CCF space. Finally, the contralateral hemisphere was assumed to be mirror-symmetric and flipped to generate a bilateral flatmap slab.

After slab generation, each voxel in CCF space was mapped to the closest voxel in flatmap space. For each voxel in the HPF and DG volumes, a 3D KDTree tree search was performed to find the closest voxel that contained a streamline point. In some cases, multiple CCF voxels mapped to the same point in flatmap space, indicating that the flatmap transformation was at a lower resolution compared to the original volume. Mappings between CCF and flatmap space were saved as a lookup table and used for mapping volumetric data. In parallel, a KDTree search was conducted in the flatmap space for each voxel to find the nearest nonzero neighbor - this table was subsequently used for nearest-neighbor interpolation for volumetric data.

### Rabies tracing and viral injections

All experiments were conducted according to the National Institutes of Health (NIH) guidelines for animal care and use and were authorized by the University of California, Irvine Institutional Animal Care and Use Committee and Institutional Biosafety Committee. To map neural circuit connectivity, mice were anesthetized with 1–2% isoflurane and positioned in a stereotaxic frame (Leica Angle Two™). Following scalp disinfection and craniotomy, viral injections were made into the dorsal CA1 (AP −1.9 mm, ML −1.4 mm, DV −1.4 mm from bregma) using a glass pipette (20–30 μm tip) and delivered by a Picospritzer at a rate of 20–30 nl/min with a 10-ms pulse duration. The helper AAVs, AAV8-DIO-TC66T-2A-GFP-2A-oG (3.8 × 10¹³ GC/ml) and AAV1-CamKII 0.4.Cre.SV40 (1.9 × 10¹³ GC/ml) were co-injected (0.1 μl). After 17 days, EnvA-RV-SADΔG-tdTomato rabies virus (CNCM, UCI) was injected at the same site (5x 108 infectious units/ml, 0.2 μl). Mice received 5 mg/kg carprofen subcutaneously to mitigate pain and inflammation and survived for 9 days to allow retrograde rabies labeling of presynaptic neurons. The mice were transcardially perfused with 20 ml of PBS and then with 40 ml of PBS containing 4% paraformaldehyde (PFA) before brain sample collection.

After perfusion, brains were processed and imaged using serial two photon tomography on a TissueCyte system with a previously published workflow (Ding et al., 2024). Cell segmentation was performed using Napari to identify putative starter cells and monosynaptic input neurons. Image volumes and cell coordinates were registered to the Allen CCFv3 template space and flatmap transformed using the CCF-to-flatmap mapping table.

### Code and data availability

All code used to generate the data, software, and figures in this text can be found at https://github.com/AllenInstitute/flatmap. Data required for creating flatmap coordinates in adult mouse can be downloaded from a public AWS S3 bucket: https://ansrs-neuroglancer-poc.s3.us-west-2.amazonaws.com/HPF+flatmap/P56_HPF.hdf5

## Acknowledgements

The authors thank the Allen Institute founder Paul G. Allen for his vision, encouragement and support, and Allen Institute chair Jody Allen. We also thank Forrest Collman and Staci Sorensen for comments on the manuscript. Research reported in this publication was supported by the National Institute On Aging of the National Institutes of Health under Award Number U01AG076791. The content is solely the responsibility of the authors and does not necessarily represent the official views of the National Institutes of Health.

---

**Figure S1.**
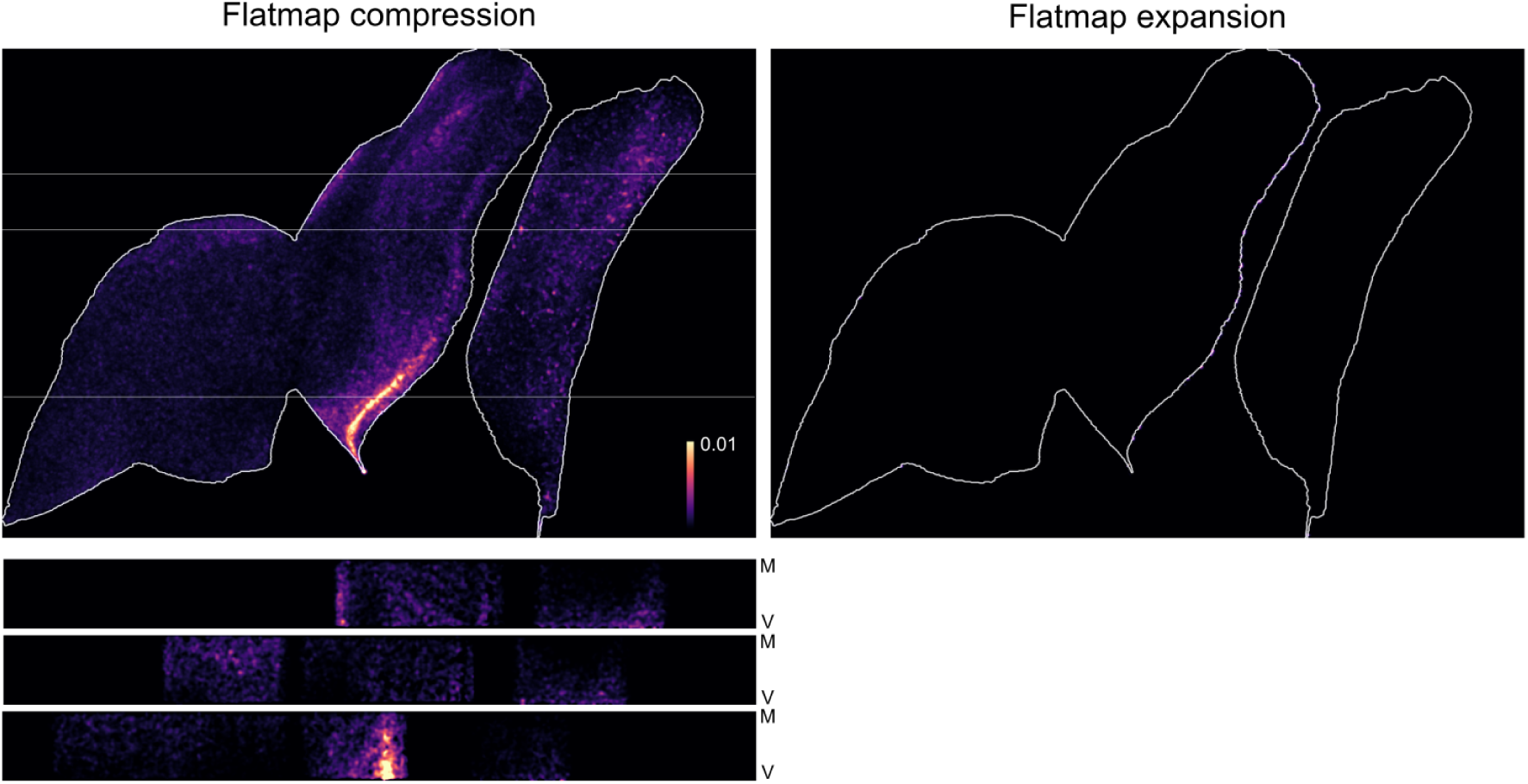
Maximum projection flatmaps showing log-transformed Jacobian determinant values, split by compressed (left) and expanded (right) regions relative to CCFv3. Heatmaps show local log-Jacobian in flatmap (top) and in cross-section along meningeal-ventricular axis (bottom). Horizontal lines show section locations.

**Figure S2.**
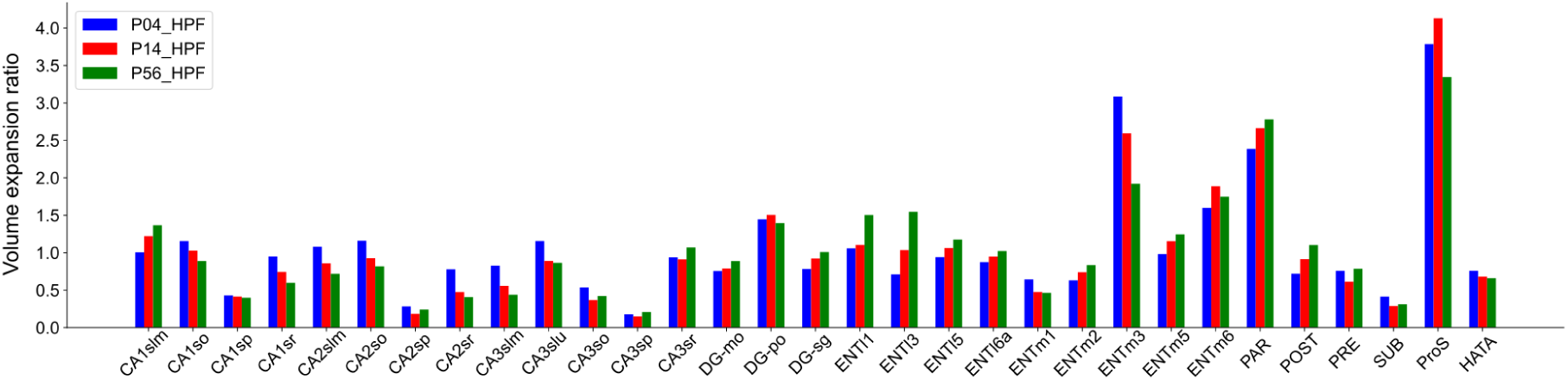
Ratio of flatmap to CCF volume for each hippocampal subregion at P4 (blue), P14 (red), and P56 (green), quantifying nonlinear deformation linked to structural curvature.

**Figure S3.**
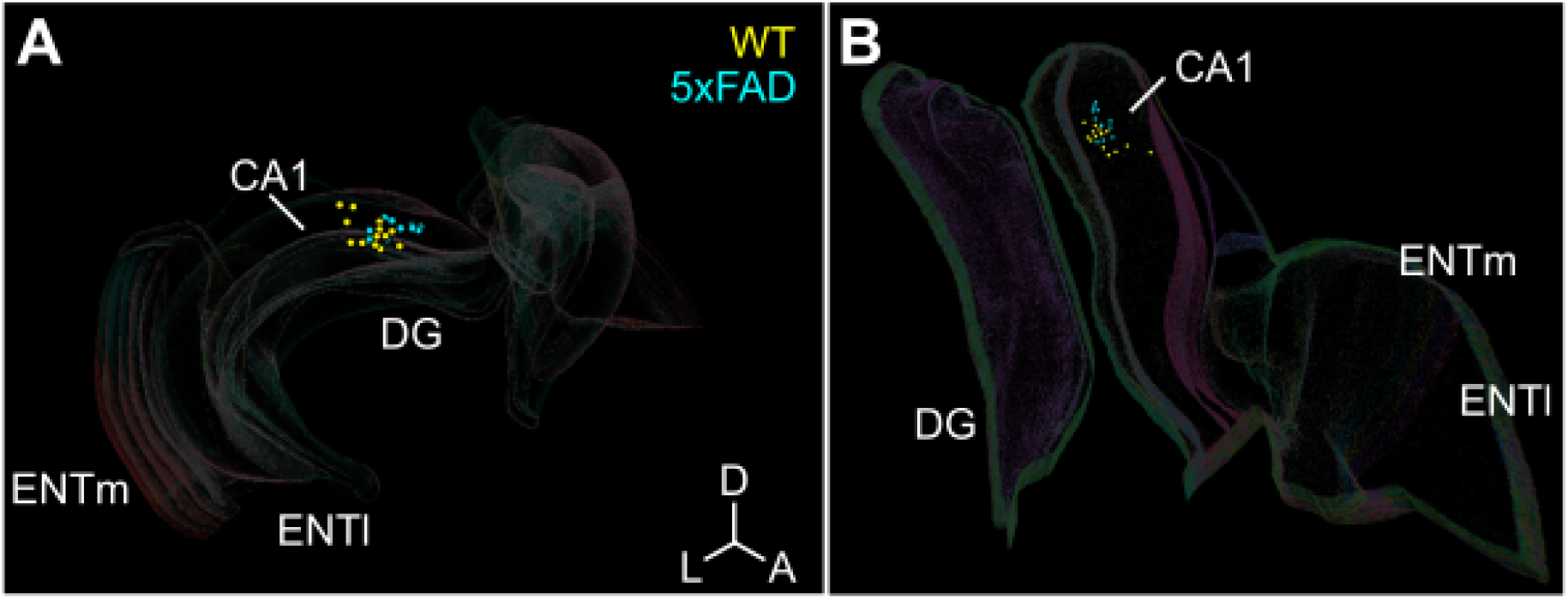
Starter cell distribution for WT (yellow) and 5xFAD (cyan) examples shown in (**A**) CCF coordinate space and (**B**) curvilinear coordinate space. Outlines show major hippocampal divisions.

## References

Amaral, D. G., & Witter, M. P. (1989). The three-dimensional organization of the hippocampal formation: a review of anatomical data. Neuroscience, 31(3), 571–591.

Angevine, J. B. (1975). Development of the Hippocampal Region. In R. L. Isaacson & K. H. Pribram (Eds.), The Hippocampus: Volume 1: Structure and Development (pp. 61–94). Springer US.

Avants, B. B., Tustison, N. J., Song, G., Cook, P. A., Klein, A., & Gee, J. C. (2011). A reproducible evaluation of ANTs similarity metric performance in brain image registration. NeuroImage, 54(3), 2033–2044.

Barry-Carroll, L., & Gomez-Nicola, D. (2024). The molecular determinants of microglial developmental dynamics. Nature Reviews. Neuroscience, 25(6), 414–427.

Barry-Carroll, L., Greulich, P., Marshall, A. R., Riecken, K., Fehse, B., Askew, K. E., Li, K., Garaschuk, O., Menassa, D. A., & Gomez-Nicola, D. (2023). Microglia colonize the developing brain by clonal expansion of highly proliferative progenitors, following allometric scaling. Cell Reports, 42(5), 112425.

Bayer, S. A. (1980). Development of the hippocampal region in the rat. II. Morphogenesis during embryonic and early postnatal life. The Journal of Comparative Neurology, 190(1), 115–134.

Bennett, H. C., Zhang, Q., Wu, Y.-T., Manjila, S. B., Chon, U., Shin, D., Vanselow, D. J., Pi, H.-J., Drew, P. J., & Kim, Y. (2024). Aging drives cerebrovascular network remodeling and functional changes in the mouse brain. Nature Communications, 15(1), 6398.

Bernhardt, B. C., Smallwood, J., Keilholz, S., & Margulies, D. S. (2022). Gradients in brain organization. NeuroImage, 251(118987), 118987.

Bolaños-Puchet, S., Teska, A., Hernando, J. B., Lu, H., Romani, A., Schürmann, F., & Reimann, M. W. (2024). Enhancement of brain atlases with laminar coordinate systems: Flatmaps and barrel column annotations. *Imaging Neuroscience (Cambridge*, Mass.), 2(imag-2-00209), 1–20.

Bolaños-Puchet, S., Teska, A., & Reimann, M. W. (2023). Enhancement of brain atlases with region-specific coordinate systems: flatmaps and barrel column annotations. In bioRxiv (p. 2023.08.24.554204). 10.1101/2023.08.24.554204

Cenquizca, L. A., & Swanson, L. W. (2007). Spatial organization of direct hippocampal field CA1 axonal projections to the rest of the cerebral cortex. Brain Research Reviews, 56(1), 1–26.

Chandrashekar, J. (2017). MouseLight Neuron Browser - Collection [Dataset]. 10.25378/janelia.c.3924088.v1

Cossart, R., & Khazipov, R. (2022). How development sculpts hippocampal circuits and function. Physiological Reviews, 102(1), 343–378.

DeKraker, J., Ferko, K. M., Lau, J. C., Köhler, S., & Khan, A. R. (2018). Unfolding the hippocampus: An intrinsic coordinate system for subfield segmentations and quantitative mapping. NeuroImage, 167, 408–418.

Diers, K., Baumeister, H., Jessen, F., Düzel, E., Berron, D., & Reuter, M. (2023). An automated, geometry-based method for hippocampal shape and thickness analysis. NeuroImage, 276, 120182.

Ding, Y., Huang, Y., Gao, P., Thai, A., Chilaparasetti, A. N., Gopi, M., Xu, X., & Li, C. (2024). Brain image data processing using collaborative data workflows on Texera. Frontiers in Neural Circuits, 18, 1398884.

Evans, A. C., Janke, A. L., Collins, D. L., & Baillet, S. (2012). Brain templates and atlases. NeuroImage, 62(2), 911–922.

Fanselow, M. S., & Dong, H.-W. (2010). Are the dorsal and ventral hippocampus functionally distinct structures? Neuron, 65(1), 7–19.

Fischl, B., Sereno, M. I., & Dale, A. M. (1999). Cortical surface-based analysis. II: Inflation, flattening, and a surface-based coordinate system. NeuroImage, 9(2), 195–207.

Frey, S., Pandya, D. N., Chakravarty, M. M., Bailey, L., Petrides, M., & Collins, D. L. (2011). An MRI based average macaque monkey stereotaxic atlas and space (MNI monkey space). NeuroImage, 55(4), 1435–1442.

Garaschuk, O., Hanse, E., & Konnerth, A. (1998). Developmental profile and synaptic origin of early network oscillations in the CA1 region of rat neonatal hippocampus. The Journal of Physiology, 507 ( *Pt 1*)(Pt 1), 219–236.

Hahn, J. D., & Duckworth, C. (2023). A brain flatmap data visualization tool for mouse, rat, and human. The Journal of Comparative Neurology, 531(10), 1008–1016.

Hahn, J. D., Swanson, L. W., Bowman, I., Foster, N. N., Zingg, B., Bienkowski, M. S., Hintiryan, H., & Dong, H.-W. (2021). An open access mouse brain flatmap and upgraded rat and human brain flatmaps based on current reference atlases. The Journal of Comparative Neurology, 529(3), 576–594.

Harris, J. A., Mihalas, S., Hirokawa, K. E., Whitesell, J. D., Choi, H., Bernard, A., Bohn, P., Caldejon, S., Casal, L., Cho, A., Feiner, A., Feng, D., Gaudreault, N., Gerfen, C. R., Graddis, N., Groblewski, P. A., Henry, A. M., Ho, A., Howard, R., … Zeng, H. (2019). Hierarchical organization of cortical and thalamic connectivity. Nature, 575(7781), 195–202.

Hawrylycz, M., Martone, M. E., Ascoli, G. A., Bjaalie, J. G., Dong, H.-W., Ghosh, S. S., Gillis, J., Hertzano, R., Haynor, D. R., Hof, P. R., Kim, Y., Lein, E., Liu, Y., Miller, J. A., Mitra, P. P., Mukamel, E., Ng, L., Osumi-Sutherland, D., Peng, H., … Zingg, B. (2023). A guide to the BRAIN Initiative Cell Census Network data ecosystem. PLoS Biology, 21(6), e3002133.

Hu, J., Li, X., Coleman, K., Schroeder, A., Ma, N., Irwin, D. J., Lee, E. B., Shinohara, R. T., & Li, M. (2021). SpaGCN: Integrating gene expression, spatial location and histology to identify spatial domains and spatially variable genes by graph convolutional network. Nature Methods, 18(11), 1342–1351.

Ishizuka, N., Weber, J., & Amaral, D. G. (1990). Organization of intrahippocampal projections originating from CA3 pyramidal cells in the rat. The Journal of Comparative Neurology, 295(4), 580–623.

Jinno, S., Fleischer, F., Eckel, S., Schmidt, V., & Kosaka, T. (2007). Spatial arrangement of microglia in the mouse hippocampus: a stereological study in comparison with astrocytes. Glia, 55(13), 1334–1347.

Jones, S. E., Buchbinder, B. R., & Aharon, I. (2000). Three-dimensional mapping of cortical thickness using Laplace’s equation. Human Brain Mapping, 11(1), 12–32.

Kaas, J. H. (1997). Topographic maps are fundamental to sensory processing. Brain Research Bulletin, 44(2), 107–112.

Kim, Y., Yang, G. R., Pradhan, K., Venkataraju, K. U., Bota, M., García Del Molino, L. C., Fitzgerald, G., Ram, K., He, M., Levine, J. M., Mitra, P., Huang, Z. J., Wang, X.-J., & Osten, P. (2017). Brain-wide maps reveal stereotyped cell-type-based cortical architecture and subcortical sexual dimorphism. Cell, 171(2), 456–469.e22.

Kleven, H., Bjerke, I. E., Clascá, F., Groenewegen, H. J., Bjaalie, J. G., & Leergaard, T. B. (2023). Waxholm Space atlas of the rat brain: a 3D atlas supporting data analysis and integration. Nature Methods. 10.1038/s41592-023-02034-3

Kleven, H., Gillespie, T. H., Zehl, L., Dickscheid, T., Bjaalie, J. G., Martone, M. E., & Leergaard, T. B. (2023). AtOM, an ontology model to standardize use of brain atlases in tools, workflows, and data infrastructures. Scientific Data, 10(1), 486.

Knierim, J. J. (2015). The hippocampus. Current Biology: CB, 25(23), R1116–R1121.

Kronman, F. N., Liwang, J. K., Betty, R., Vanselow, D. J., Wu, Y.-T., Tustison, N. J., Bhandiwad, A., Manjila, S. B., Minteer, J. A., Shin, D., Lee, C. H., Patil, R., Duda, J. T., Xue, J., Lin, Y., Cheng, K. C., Puelles, L., Gee, J. C., Zhang, J., … Kim, Y. (2024). Developmental mouse brain common coordinate framework. Nature Communications, 15(1), 9072.

Leergaard, T. B., & Bjaalie, J. G. (2022). Atlas-based data integration for mapping the connections and architecture of the brain. Science (New York, N.Y.), 378(6619), 488–492.

Liu, L., Yun, Z., Manubens-Gil, L., Chen, H., Xiong, F., Dong, H., Zeng, H., Hawrylycz, M., Ascoli, G. A., & Peng, H. (2025). Connectivity of single neurons classifies cell subtypes in mouse brains. Nature Methods, 22(4), 861–873.

Liwang, J. K., Kronman, F. N., Pi, H.-J., Wu, Y.-T., Vanselow, D. J., Manjila, S. B., Parmaksiz, D., Shin, D., Ben-Simon, Y., Taormina, M., Way, S. W., Zeng, H., Tasic, B., Ng, L., & Kim, Y. (2025). epDevAtlas: mapping GABAergic cells and microglia in the early postnatal mouse brain. Nature Communications, 16(1), 9538.

Li, X. G., Somogyi, P., Ylinen, A., & Buzsáki, G. (1994). The hippocampal CA3 network: an in vivo intracellular labeling study: THE HIPPOCAMPAL CA3 NETWORK. The Journal of Comparative Neurology, 339(2), 181–208.

Milnor, J. (1969). A problem in cartography. The American Mathematical Monthly: The Official Journal of the Mathematical Association of America, 76(10), 1101–1112.

Ng, L., Lau, C., Sunkin, S. M., Bernard, A., Chakravarty, M. M., Lein, E. S., Jones, A. R., & Hawrylycz, M. (2010). Surface-based mapping of gene expression and probabilistic expression maps in the mouse cortex. Methods, 50(2), 55–62.

Oh, S. W., Harris, J. A., Ng, L., Winslow, B., Cain, N., Mihalas, S., Wang, Q., Lau, C., Kuan, L., Henry, A. M., Mortrud, M. T., Ouellette, B., Nguyen, T. N., Sorensen, S. A., Slaughterbeck, C. R., Wakeman, W., Li, Y., Feng, D., Ho, A., … Zeng, H. (2014). A mesoscale connectome of the mouse brain. Nature, 508(7495), 207–214.

Olsen, L. C., Galler, M., Witter, M. P., Saetrom, P., & O’Reilly, K. C. (2023). Transcriptional development of the hippocampus and the dorsal-intermediate-ventral axis in rats. Hippocampus, 33(9), 1028–1047.

Paolicelli, R. C., Bolasco, G., Pagani, F., Maggi, L., Scianni, M., Panzanelli, P., Giustetto, M., Ferreira, T. A., Guiducci, E., Dumas, L., Ragozzino, D., & Gross, C. T. (2011). Synaptic pruning by microglia is necessary for normal brain development. Science (New York, N.Y.), 333(6048), 1456–1458.

Peng, H., Xie, P., Liu, L., Kuang, X., Wang, Y., Qu, L., Gong, H., Jiang, S., Li, A., Ruan, Z., Ding, L., Yao, Z., Chen, C., Chen, M., Daigle, T. L., Dalley, R., Ding, Z., Duan, Y., Feiner, A., … Zeng, H. (2021). Morphological diversity of single neurons in molecularly defined cell types. Nature, 598(7879), 174–181.

Puelles, L., Alonso, A., García-Calero, E., & Martínez-de-la-Torre, M. (2019). Concentric ring topology of mammalian cortical sectors and relevance for patterning studies. The Journal of Comparative Neurology, 527(10), 1731–1752.

Puelles, L., Harrison, M., Paxinos, G., & Watson, C. (2013). A developmental ontology for the mammalian brain based on the prosomeric model. Trends in Neurosciences, 36(10), 570–578.

Qiu, S., Hu, Y., Huang, Y., Gao, T., Wang, X., Wang, D., Ren, B., Shi, X., Chen, Y., Wang, X., Wang, D., Han, L., Liang, Y., Liu, D., Liu, Q., Deng, L., Chen, Z., Zhan, L., Chen, T., … Xu, C. (2024). Whole-brain spatial organization of hippocampal single-neuron projectomes. Science (New York, N.Y.), 383(6682), eadj9198.

Sherk, H. (1992). Flattening the cerebral cortex by computer. Journal of Neuroscience Methods, 41(3), 255–267.

Strange, B. A., Witter, M. P., Lein, E. S., & Moser, E. I. (2014). Functional organization of the hippocampal longitudinal axis. Nature Reviews. Neuroscience, 15(10), 655–669.

Swanson, L. W., & Hahn, J. D. (2020). A qualitative solution with quantitative potential for the mouse hippocampal cortex flatmap problem. Proceedings of the National Academy of Sciences of the United States of America, 117(6), 3220–3231.

Swanson, L. W., Wyss, J. M., & Cowan, W. M. (1978). An autoradiographic study of the organization of intrahippocampal association pathways in the rat. The Journal of Comparative Neurology, 181(4), 681–715.

Tenenbaum, J. B., de Silva, V., & Langford, J. C. (2000). A global geometric framework for nonlinear dimensionality reduction. Science, 290(5500), 2319–2323.

Thompson, C. L., Ng, L., Menon, V., Martinez, S., Lee, C.-K., Glattfelder, K., Sunkin, S. M., Henry, A., Lau, C., Dang, C., Garcia-Lopez, R., Martinez-Ferre, A., Pombero, A., Rubenstein, J. L. R., Wakeman, W. B., Hohmann, J., Dee, N., Sodt, A. J., Young, R., … Jones, A. R. (2014). A high-resolution spatiotemporal atlas of gene expression of the developing mouse brain. Neuron, 83(2), 309–323.

Townes, F. W., & Engelhardt, B. E. (2023). Nonnegative spatial factorization applied to spatial genomics. Nature Methods, 20(2), 229–238.

Van Essen, D. C., & Maunsell, J. H. (1980). Two-dimensional maps of the cerebral cortex. The Journal of Comparative Neurology, 191(2), 255–281.

van Groen, T., Miettinen, P., & Kadish, I. (2003). The entorhinal cortex of the mouse: organization of the projection to the hippocampal formation. Hippocampus, 13(1), 133–149.

Wang, Q., Ding, S.-L., Li, Y., Royall, J., Feng, D., Lesnar, P., Graddis, N., Naeemi, M., Facer, B., Ho, A., Dolbeare, T., Blanchard, B., Dee, N., Wakeman, W., Hirokawa, K. E., Szafer, A., Sunkin, S. M., Oh, S. W., Bernard, A., … Ng, L. (2020). The Allen Mouse Brain Common Coordinate Framework: A 3D Reference Atlas. Cell, 181(4), 936–953.e20.

Witter, M. P., Groenewegen, H. J., Lopes da Silva, F. H., & Lohman, A. H. (1989). Functional organization of the extrinsic and intrinsic circuitry of the parahippocampal region. Progress in Neurobiology, 33(3), 161–253.

Wong, T., Zhang, X. L., Asl, M. N., Wu, C. P., Carlen, P. L., & Zhang, L. (2005). Postnatal development of intrinsic GABAergic rhythms in mouse hippocampus. Neuroscience, 134(1), 107–120.

Wu, Y.-T., Bennett, H. C., Chon, U., Vanselow, D. J., Zhang, Q., Muñoz-Castañeda, R., Cheng, K. C., Osten, P., Drew, P. J., & Kim, Y. (2022). Quantitative relationship between cerebrovascular network and neuronal cell types in mice. Cell Reports, 39(12), 110978.

Yao, Z., van Velthoven, C. T. J., Kunst, M., Zhang, M., McMillen, D., Lee, C., Jung, W., Goldy, J., Abdelhak, A., Aitken, M., Baker, K., Baker, P., Barkan, E., Bertagnolli, D., Bhandiwad, A., Bielstein, C., Bishwakarma, P., Campos, J., Carey, D., … Zeng, H. (2023). A high-resolution transcriptomic and spatial atlas of cell types in the whole mouse brain. Nature, 624(7991), 317–332.

Zhang, M., Pan, X., Jung, W., Halpern, A. R., Eichhorn, S. W., Lei, Z., Cohen, L., Smith, K. A., Tasic, B., Yao, Z., Zeng, H., & Zhuang, X. (2023). Molecularly defined and spatially resolved cell atlas of the whole mouse brain. Nature, 624(7991), 343–354.

Zhao, K., Wang, D., Wang, D., Chen, P., Wei, Y., Tu, L., Chen, Y., Tang, Y., Yao, H., Zhou, B., Lu, J., Wang, P., Liao, Z., Chen, Y., Han, Y., Zhang, X., & Liu, Y. (2024). Macroscale connectome topographical structure reveals the biomechanisms of brain dysfunction in Alzheimer’s disease. Science Advances, 10(41), eado8837.

